# Identification of KIFC1 as a putative vulnerability in lung cancers with centrosome amplification

**DOI:** 10.1101/2024.01.09.574105

**Authors:** Christopher Zhang, Benson Wu, Yin Fang Wu, Caterina di Ciano-Oliveira, Isabel Soria-Bretones, Nhu-An Pham, Andrew J. Elia, Raj Chari, Wan L. Lam, Mark R. Bray, Tak W. Mak, Ming-Sound Tsao, David W. Cescon, Kelsie L. Thu

**Affiliations:** Department of Laboratory Medicine and Pathobiology, University of Toronto; Keenan Research Centre for Biomedical Science, St. Michael’s Hospital, Unity Health Toronto; Princess Margaret Cancer Centre, University Health Network; Laboratory Animal Sciences Program, Frederick National Laboratory for Cancer Research; Department of Integrative Oncology, BC Cancer Research Institute; Department of Pathology and Laboratory Medicine, University of British Columbia; Departments of Immunology University of Toronto; Departments of Medical Biophysics, University of Toronto; Institute of Medical Science, University of Toronto

## Abstract

Centrosome amplification (CA), an abnormal increase in the number of centrosomes in the cell, is a recurrent phenomenon in lung and other malignancies. Although CA contributes to tumor development and progression by promoting genomic instability (GIN), it also induces mitotic stress that jeopardizes cellular integrity. The presence of extra centrosomes leads to the formation of multipolar mitotic spindles prone to causing lethal chromosome segregation errors during cell division. To sustain the benefits of CA, malignant cells are dependent on adaptive mechanisms to mitigate its detrimental consequences, and these mechanisms represent therapeutic vulnerabilities. We aimed to discover genetic dependencies associated with CA in lung cancer. Combining a CRISPR/Cas9 functional genomics screen with analyses of tumor genomic data, we identified the motor protein KIFC1 as a putative vulnerability specifically in lung cancers with CA. KIFC1 expression was positively correlated with CA in lung adenocarcinoma (LUAD) cell lines and with a gene expression signature predictive of CA in LUAD tumor tissues. High *KIFC1* expression was associated with worse patient outcomes, smoking history, and indicators of GIN. KIFC1 loss-of-function sensitized LUAD cells to potentiation of CA and sensitization was associated with a diminished ability of KIFC1-depleted cells to cluster extra centrosomes into pseudo-bipolar mitotic spindles. Our work suggests that KIFC1 inhibition represents a novel approach for potentiating GIN to lethal levels in LC with CA by forcing cells to divide with multipolar spindles, rationalizing the clinical development of KIFC1 inhibitors and further studies to investigate its therapeutic potential.

## INTRODUCTION

Lung cancer (LC) is the leading cause of cancer death worldwide[1] and is broadly divided into small-cell lung cancer (SCLC) and non-small cell lung cancer (NSCLC). Of the various NSCLC subtypes, lung adenocarcinoma (LUAD) is most prevalent, accounting for ∼40% of all cases[2]. Although advances in targeted therapies and immunotherapy have expanded the treatment landscape for LC[3], the 5-year net survival rate has not risen above 22%[4]. This highlights the need for new therapeutic approaches that target mechanisms supporting lung tumor growth and progression.

Centrosomes are microtubule-organizing organelles with important roles in cell motility and polarity, and generation of mitotic spindles required to segregate chromosomes during mitosis[5]. Centrosome amplification (CA), an abnormal increase in the number of centrosomes in a cell, is a common feature of many malignancies including LC[5–7]. Several routes to CA have been described including DNA damage, failure of cytokinesis, centriole overduplication, and carcinogen exposure[8–11]. The presence of extra centrosomes has been shown to promote genomic instability (GIN)[12, 13] which fuels tumor adaptation to stress such as that induced by anti-cancer therapy[14]. This fitness advantage may explain the prevalence of CA in cancer; however, having supernumerary centrosomes comes with a cost. Cells with CA are prone to forming multipolar mitotic spindles that if left uncorrected cause severe chromosome segregation errors, mitotic catastrophe, and cell death[15]. Thus, cancer cells must manage extra centrosomes to prevent multipolar divisions and their deleterious effects. Several coping mechanisms have been proposed with clustering of extra centrosomes into pseudo-bipolar spindles being the most well established[5, 12, 16]. As such, the proteins regulating these adaptive mechanisms represent vulnerabilities and potential therapeutic targets in cancer cells with CA.

Multiple reports have demonstrated the occurrence and clinical relevance of CA in LC[6, 7, 17]. However, despite its occurrence, the mechanisms LC use to cope with CA have not been directly investigated. We sought to identify collateral dependencies associated with the presence of extra centrosomes in LC models. Integrating a CRISPR/Cas9 screen with tumor genomic analyses, we identified KIFC1 as a putative survival factor for LUAD with CA. We discovered that KIFC1 inactivation is lethal specifically in LC cells with CA. Our analyses of cell lines and independent tumor datasets revealed that KIFC1 expression is positively correlated with CA in LUAD suggesting centrosome number could indicate LC sensitivity to KIFC1 inhibition. We found that KIFC1 depletion sensitized LUAD cells to potentiation of CA and that this phenotype was associated with a diminished ability to cluster extra centrosomes. Collectively, these findings nominate KIFC1 as a therapeutic target in LUAD with CA, rationalizing the development of KIFC1-specific inhibitors and further preclinical investigation of its therapeutic potential.

## METHODS

### Tumor and cell line genomic analyses

RNA-sequencing (RNASeqV2) and associated clinical data for The Cancer Genome Atlas (TCGA) Pan-Cancer (PANCAN) cohort were accessed from the National Cancer Institute’s Genomic Data Commons portal[18]. Mutation status for the most common oncogenic drivers (*EGFR, KRAS, MET, ROS1, RET, ALK, NRAS and BRAF*) in TCGA LUAD tumors was obtained from cBioPortal[19]. Gene expression and essentiality data for the Cancer Cell Line Encyclopedia (CCLE) were obtained from the DepMap database (https://depmap.org/portal/)[20]. Normalized microarray-derived gene expression data for 4 additional LUAD cohorts were obtained from the Gene Expression Omnibus (GSE50081, GSE31210, GSE68465, GSE75037). CA20 gene expression signature scores for TCGA lung tumors and CCLE cell lines were obtained from a published study[21], and were calculated for additional cohorts as described[22].

### Immunohistochemistry

Tumor tissues were banked after informed written patient consent following a protocol approved by the University Health Network Research Ethics Board. Formalin-fixed paraffin embedded tissues from 10 NSCLC patient-derived xenograft (PDX) models were processed for immunohistochemistry to detect centrosomes using the anti-pericentrin antibody ab4448 (Abcam, 1:1000 dilution). Mitotic cells were classified as having CA if they contained more than 2 centrosomes and the proportion with CA was calculated for each tumor. At least 30 mitotic cells were counted per tumor tissue section.

### Cell culture and lentivirus production

Cell lines were grown at 37°C and 5% CO2 in DMEM, Hams F12 or RPMI 1640 media supplemented with 10% fetal bovine serum, 2mM glutamine, and 100U/ml of penicillin-streptomycin. NCI-H1299, - H1975, -H1944, -H2291, -H1792, -H2122, -H23, -H2030, -H1568, -H1650, HCC827, HCC2935 were generous gifts from the late Dr. Adi Gazdar. PC9 was gifted from Dr. William Lockwood and A549 was gifted from Dr. Haibo Zhang. Cells were confirmed to be negative for mycoplasma using a PCR-based assay[23]. Platinum-E (PE) cells were obtained from Cell BioLabs and used to produce lentivirus by transfection with psPAX2 and pMD2.G (Addgene #12260, #12259) using Lipofectamine 3000.

### CRISPR/Cas9 genetic screen

H1299 cells were engineered with stable Cas9 expression using Lenti-Cas9-blast (Addgene #52962). Efficient nuclease activity was confirmed using a reporter system (Addgene #67979, #67980; **Supp. Fig. 1**). We synthesized a custom single guide RNA (sgRNA) knockout (KO) library with 13,243 sgRNA targeting 3,319 genes that are “druggable” with FDA-approved or investigational compounds annotated in the DrugBank and bindingDB databases[24, 25], as well as negative controls (sgLacZ, sgLuciferase, sgGFP) (**Supp. Table 1**). This library was cloned into pRC0162 (Addgene #195319) and sequence verified. H1299-Cas9 cells were infected with the library at a multiplicity of infection of 30% for single integrations per cell to yield 500X library coverage. Following 48hr of puromycin selection, cells were pooled and split into 3 replicates each for treatment with vehicle (DMSO) or 7.5nM of CFI-400945 (945). Ten million cells were collected at time zero (T0) and the remainder were cultured in the presence of DMSO or 945 for 17 cell doublings and then harvested for targeted sgRNA sequencing. Screens hits were identified using the DrugZ algorithm[26].

**Figure 1.**
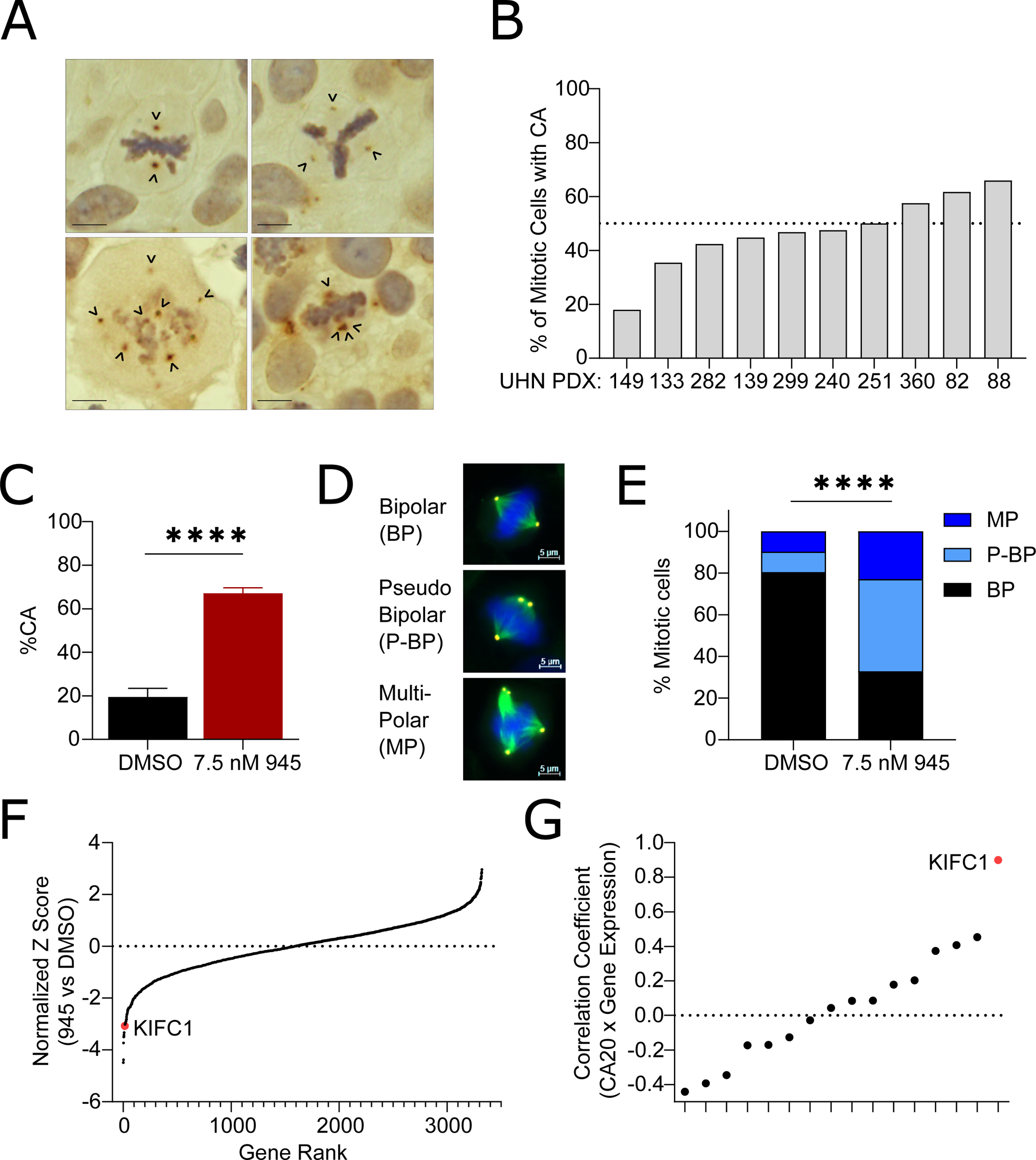
CRISPR/Cas9 genetic screen identifies KIFC1 as a dependency in lung cancers with centrosome amplification (CA). (A) Representative IHC images of mitotic cells in NSCLC tissues from patient-derived xenografts (PDX) stained for the centrosome marker pericentrin. Arrowheads point to centrosomes. Scale bar indicates 10um. (B) Characterization of CA in 10 NSCLC PDX. Centrosomes were scored in mitotic cells only and cells were classified as having CA if they contained > 2 pericentrin foci. (C) Potentiation of CA in H1299-Cas9+ cells by 945 treatment during the CRISPR screen. Cells were sampled from each replicate (N=3/treatment arm). Bar plots indicate mean with SD, asterisks indicate t-test p<0.0001. (D) Representative IF images exhibiting bipolar (BP), multipolar (MP), and pseudo-bipolar (P-BP) mitotic spindles in H1299 cells. Scale bar indicates 5um. (E) Characterization of mitotic spindles in H1299-Cas9+ cells sampled during the CRISPR screen. Counts were pooled across the 3 replicates for each treatment condition. Asterisks indicate chi-square test p<0.0001. A minimum of 160 total mitotic cells were counted for each treatment condition for IF analyses in (C) and (E). (F) Normalized Z-scores for each gene targeted in the sgRNA library calculated with the DrugZ algorithm. Z-scores represent the relative abundance of sgRNAs in cells treated with 945 relative to DMSO control. Negative Z-scores indicate genes targeted by sgRNAs that were significantly depleted in cells in which CA was potentiated by 945. (G) Correlations between gene expression and CA20 scores for the top 16 genes identified by the CRISPR screen in the TCGA LUAD cohort. Correlation coefficients indicate those for Pearson’s correlation analyses.

### Competition Assays

H1975, PC9, and A549 were engineered to express functional Cas9 as above. sgRNA targeting AAVS1 (AV, negative control) or KIFC1 (K3, K4) were cloned into LentiGuide-puro-NLS-GFP or LentiGuide-puro-NLS-mCherry (#185473, 185474) and transduced into Cas9+ cells. sgRNA sequences are provided in **Supp. Table 2**. mCherry+ AV (wildtype, WT) were mixed 1:1 with GFP+ K3/K4/AV cells to compare the relative fitness of cells with and without KIFC1 perturbation. Cells were treated with DMSO or 7.5nM 945 and mCherry/GFP+ fractions were measured by flow cytometry over multiple passages.

### KIFC1 knockdown, dose response, and viability assays

Human Silencer Select siRNA assays s7906, s7907, and Negative Control No. 1 (Thermo Fisher Scientific) were used to knockdown (KD) KIFC1. Cells were transfected with siRNA using Lipofectamine RNAiMAX (50nM in H1299, H1975, A549; 75nM in PC9) and 24 hours later, were sampled to confirm KD and plated for functional experiments. Dose response, clonogenic survival, and AZ82 cytotoxicity assays were done using sulforhodamine B staining as previously described[27]. Proliferation was assessed using the ATP-Lite assay.

### Immunofluorescence, immunoblotting, and real-time quantitative PCR

Immunofluorescence (IF) was used to detect centrosomes using the following antibodies: CEP192 (1:1000, Bethyl Labs#A302-324A)[28], ɑ-tubulin (1:1000, Sigma-Aldrich #T9026), goat anti-mouse AlexaFluor-488, and goat anti-rabbit AlexaFluor-555. DNA was stained with DAPI. CA was calculated as described for IHC above. Western blots were done using the following antibodies: KIFC1 (1:1000, CST#12313), GAPDH (1:5000, CST#97166), and β-ACTIN (1:5000, Proteintech#66009-1-IG). RT-qPCR to confirm siRNA-mediated *KIFC1* knockdown was done using *HPRT1* as the endogenous control. Primer sequences are listed in **Supp. Table 2**.

### Statistics

Statistical analyses are described in the figure legends and were performed using R and GraphPad Prism.

## RESULTS

### Centrosome amplification (CA) is prevalent in NSCLC tissues

Previous studies have reported centrosome abnormalities in 29-53% of NSCLC tissues[6, 7]. We evaluated CA in 10 patient-derived xenograft (PDX) models of NSCLC (4 LUAD, 3 squamous cell carcinoma (LUSC), and 3 large cell carcinoma) using IHC to detect the established centrosome marker, pericentrin[29]. Cases were chosen to represent different NSCLC subtypes. Our assessment of these tumors confirmed that CA is prevalent in NSCLC **(Fig. 1A)**. CA was detected in mitotic cells from all 10 PDX, with the percentage of cells with CA ranging from 18-66% **(Fig. 1B)**. We observed that centrosomes in cells with CA were disorganized in multipolar spindles or clustered into pseudo-bipolar spindles[12, 16] **(Fig. 1A)**. The presence of CA in clinical NSCLC tissues rationalized investigation of vulnerabilities associated with extra centrosomes in cell models.

### CRISPR screen identifies KIFC1 as a putative CA-coping factor in LC

To discover genetic dependencies in LC cells with CA, we performed a CRISPR/Cas9 screen in the LUAD cell line, H1299. We selected H1299 because of its high Cas9 editing efficiency (**Supp.** Fig. 1A), its relatively high basal CA which implied adaptation to CA (**Fig.1C; Supp. Fig. 4C**), and its amenability to potentiation of CA with low doses of the PLK4 inhibitor, CFI-400945 (945)[30] (**Supp. Fig. 1B)**. H1299-Cas9 cells were infected with a custom sgRNA KO library targeting ∼3,300 “druggable” genes, grown in the presence of DMSO or sublethal dose of 945 (7.5nM) that potentiated CA for 17 cell doublings (**Supp. Fig. 1C**), and then harvested for sgRNA sequencing. We first compared sgRNA abundance in DMSO-treated cells at end point to cells harvested at screen onset (T0) to confirm the depletion of sgRNA targeting essential genes. This identified *MYC* as the top hit with 7 other genes classified by the DepMap as “common essential genes” ranking in the top 10 hits (*CDC25, SRSF1, RPL4, PSMA3, RPL14, ATP2A2,* and *SCD*)[20] **(Supp. Fig. 1D-E)**. Notably, *NRAS*, an established oncogenic driver in H1299[31] was identified as the 10th ranked hit **(Supp. Fig. 1D-E)**. Detection of these essential genes confirmed the technical success of our CRISPR/Cas9 screen.

Sampling of H1299 cells during the screen confirmed that 7.5nM 945 potentiated CA to ∼65% of cells and increased the incidence of multipolar spindles compared to DMSO (**Fig.1C-E**). Accordingly, growth in the presence of 945 was slower (**Supp. Fig. 1C**). We reasoned that cells containing sgRNA targeting genes essential for survival in the presence of extra centrosomes would be selectively depleted in populations treated with 945 compared to DMSO. Analysis of sgRNA abundance in 945-versus DMSO-treated populations identified 16 putative CA survival genes with an FDR q-value < 0.25 (**Fig.1F; Supp.Table 3**). To prioritize hits from our screen, we examined their correlations with a gene expression signature of CA called CA20 in the TCGA LUAD cohort[32]. CA20 is a composite of 20 genes associated with the presence of CA that was validated in clinical breast tumors[21, 22]. We hypothesized that genes important for coping with extra centrosomes would be highly expressed in tumors with CA and positively correlated with CA20 scores. Of the 16 candidates, none were CA20 signature genes and KIFC1 exhibited the strongest positive correlation with CA20 in LUAD (**Fig.1G**). KIFC1 is a minus-end directed kinesin motor protein involved in mitotic spindle assembly and organization, as well as vesicle and organelle trafficking[33]. It also has a well-described role in centrosome clustering in cancer cells[33]. Thus, our CA sensitization screen suggested that H1299 cells are dependent on KIFC1 to cope with CA, nominating KIFC1 as a therapeutic vulnerability in LC with supernumerary centrosomes.

### KIFC1 is associated with clinical factors in LUAD

We next aimed to determine the clinical relevance of KIFC1 in patient-derived tumor tissues by analyzing KIFC1 mRNA expression in the TCGA LUAD cohort. Consistent with previous reports in pan-cancer or LC-focused studies [34–37], KIFC1 was significantly overexpressed in LUAD compared to non-malignant lung tissues (**Fig.2A**), and LUAD patients whose tumors had high KIFC1 expression had significantly worse 5 year survival outcomes than those with low expression in a univariate model (**Fig.2B**). KIFC1 expression remained significantly associated with overall survival in a Cox-proportional hazards model including additional prognostic factors including age, stage, sex and smoking history **(Table 1)**. Expression was also elevated in tumors of greater stage, and for the first time, we discovered that expression was significantly higher in smokers relative to non-smokers (**Fig.2C-D**). Lastly, we found that KIFC1 expression was modestly elevated in males relative to females and in tumors without oncogenic driver mutations compared to those with oncogenic alterations (**Supp. Fig. 2A**). An ANOVA suggested that smoking history was the clinical factor most strongly associated with KIFC1 expression **(Table 2)**. Although KIFC1 was overexpressed and correlated with CA20 in LUSC, we did not observe any significant associations between KIFC1 expression and patient characteristics or survival (**Supp. Fig. 3**). We then investigated the robustness of these associations in 4 independent LUAD cohorts with gene expression and clinical data available (GSE75037, GSE50081, GSE31210, GSE68465) (**Supp. Fig. 2B-F**)[38–41]. These analyses confirmed that high KIFC1 expression was associated with smoking (3/4 cohorts), advanced stage (4/4 cohorts), and worse 5-year survival (4/4 cohorts). Together these findings indicate the clinical and prognostic significance of KIFC1 in LUAD.

**Figure 2.**
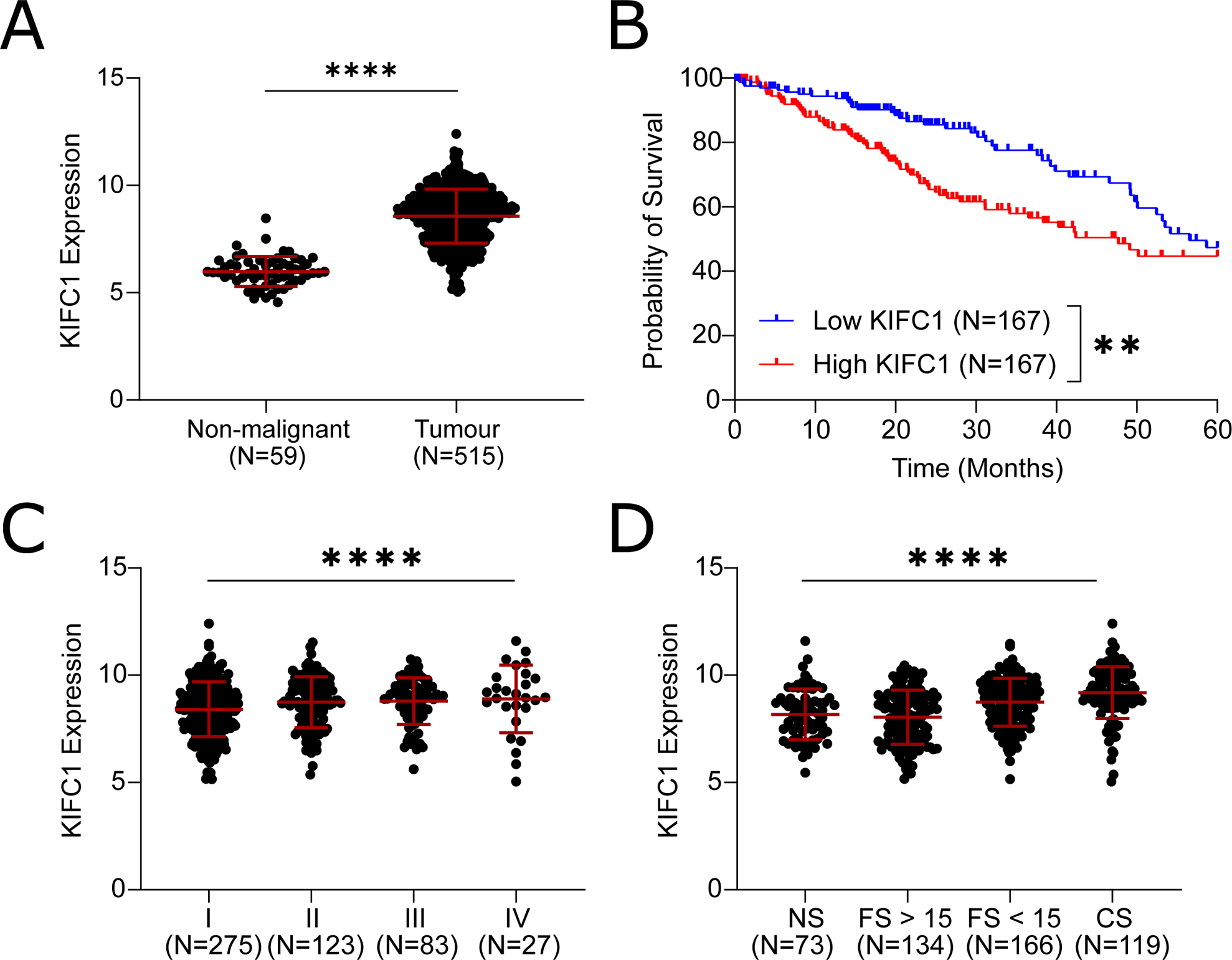
KIFC1 expression is associated with multiple clinical factors in LUAD patients. KIFC1 expression was analyzed in the TCGA LUAD cohort to investigate associations with malignancy (A), 5-year survival (B), stage (C), and smoking history (D). Asterisks indicate significance for Mann Whitney or Kruskal-Wallis tests (*p<0.05, **p<0.01, ****p<0.0001). Error bars indicate mean and SD. NS = never smokers, CS = current smokers, FS < 15 = former smoker for fewer than 15 years, FS > 15 = former smoker for over 15 years. The association between KIFC1 expression and outcome was assessed in patients whose tumors exhibited KIFC1 expression ranking in the top and bottom tertiles of expression. Asterisks indicate p<0.01 for a log-rank test on the Kaplan-Meier survival curves.

### CA and GIN correlate with KIFC1 expression in LUAD models and tissues

Since our screen nominated KIFC1 as a genetic dependency of LC cells with CA, we suspected KIFC1 expression would be higher in cells with extra centrosomes. To test this hypothesis, we evaluated KIFC1 protein expression and CA in a panel of LUAD cell lines. Both KIFC1 expression and CA were variable across models (**Supp. Fig. 4A-C**). In support of our hypothesis, we identified a positive correlation between KIFC1 protein expression and CA across LUAD cell lines (**Fig.3A**). We next evaluated the association between CA20 and KIFC1 dependency scores from the DepMap database in a larger panel of LC cell lines from the Cancer Cell Line Encyclopedia (CCLE) collection[20]. Our analyses revealed no association between CA20 and KIFC1 essentiality across 52 NSCLC or 13 LUSC lines, but indicated trends towards a greater dependency on KIFC1 in 15 LUAD and 11 small-cell lung cancer cell lines predicted to have high levels of CA (**Fig.3B-C; Supp. Fig. 4D-E**). Thus, KIFC1 may be a LUAD-specific dependency. As noted above (**Fig.1G**), we observed a strong positive correlation between KIFC1 expression and CA20 in LUAD tumors from the TCGA (**Fig.3D**), and we validated this correlation in 4 independent cohorts (**Supp. Fig. 2E**). Because CA drives genomic instability (GIN)[13, 42, 43], we also investigated associations between KIFC1 expression and measures of GIN in the TCGA LUAD cohort. This revealed significant positive correlations between KIFC1 and fraction of genome altered and aneuploidy scores[18] (**Fig.3E-F**). Collectively, these findings demonstrate that KIFC1 expression is associated with CA and GIN in LUAD models and tissues.

**Figure 3.**
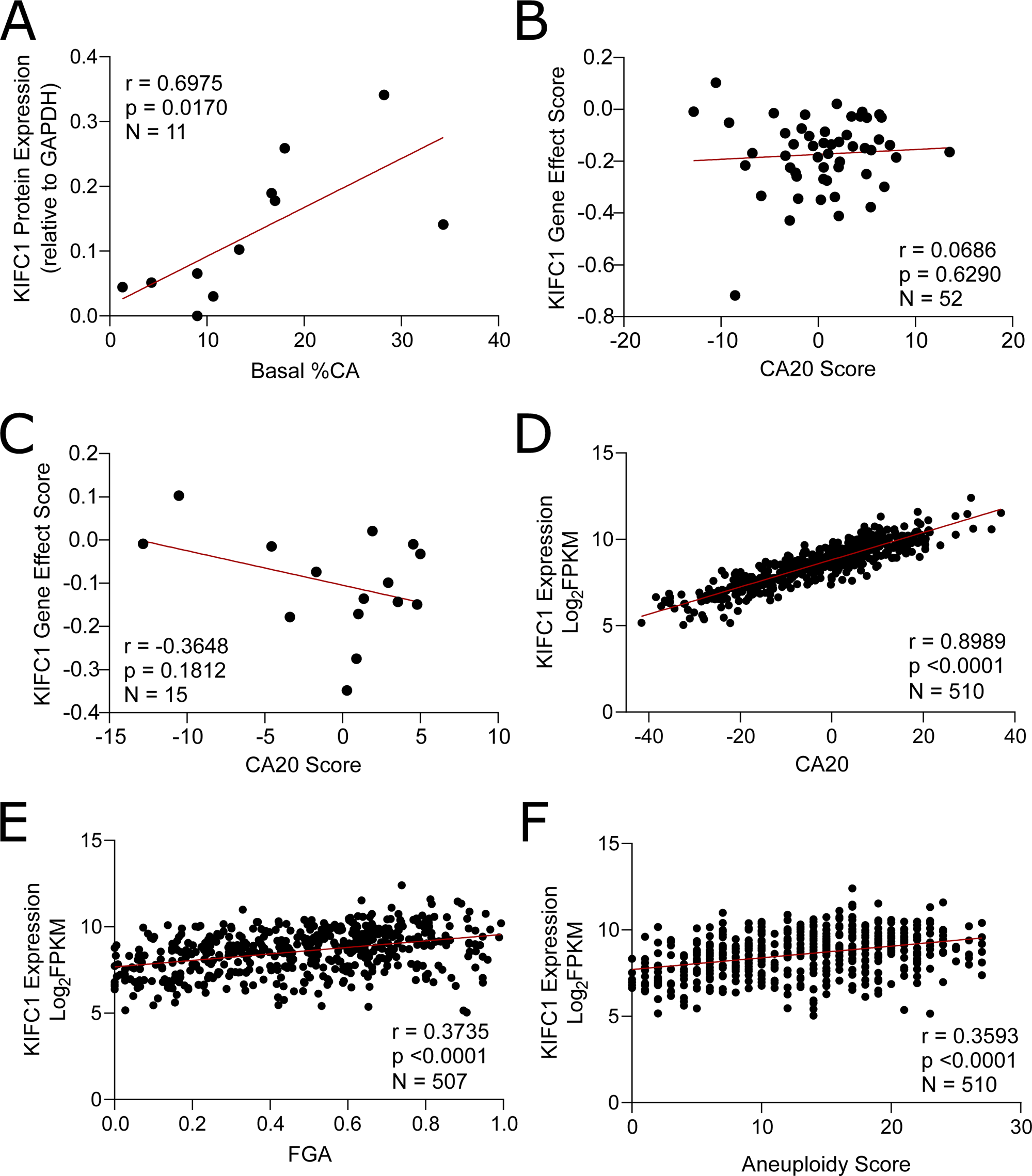
Centrosome amplification and genomic instability are positively correlated with KIFC1 expression in LUAD models and tissues. (A) Correlation between KIFC1 expression and CA in 11 LUAD models. Protein expression was quantified from western blots and CA from IF conducted under basal conditions (Supp. Fig. 4). (B,C) Correlation between KIFC1 dependency and CA20 in NSCLC (B) and LUAD (C) cell lines from the CCLE. KIFC1 gene effect scores were accessed from the DepMap database and negative scores indicate genetic dependency. CA20 scores were obtained from the de Almeida 2019 study. (D) Association between KIFC1 mRNA expression and CA20 in the TCGA LUAD cohort. (E,F) Correlation between KIFC1 mRNA expression and fraction of genome altered (FGA) and aneuploidy score. FGA and aneuploidy scores were obtained from cBioPortal. All correlation coefficients indicated were calculated using Pearson’s correlation analyses.

### LUAD cells are dependent on KIFC1 for survival upon CA potentiation

We next sought to functionally validate KIFC1 as a dependency of LC cells with CA using *in vitro* growth competition assays that mimic CRISPR screen conditions. We used models with different endogenous levels of KIFC1 and CA and examined the effects of KIFC1 loss-of-function (LOF) on cellular fitness. H1299 and H1975 were selected as representative models with relatively high basal KIFC1 expression and CA, and A549 and PC9 as models with low KIFC1 and CA (**Supp. Fig. 4**). Treatment with low-dose 945 potentiated CA and increased the proportion of multipolar spindles in each LUAD line except A549 (**Fig.4A-B**). Cas9+ derivatives of these lines were transduced with a GFP+ lentivector encoding independent sgRNA targeting KIFC1 (K3 or K4), or a GFP+ or mCherry+ lentivector encoding an sgRNA targeting AAVS1 (AV, wildtype (WT) control). CRISPR-mediated KIFC1 inactivation was evident in all models relative to controls (**Fig.4C**). We competed GFP+ WT or GFP+ K3/K4 cells with mCherry+ WT cells in the presence of DMSO or 945 and monitored the GFP:mCherry+ ratio over several passages. These experiments validated KIFC1 as a hit from our screen, as GFP+ cells were depleted in the 945-treatment condition over time (**Fig.4D**), confirming that KIFC1 LOF sensitized H1299 cells to CA potentiation. We observed the same phenotype in H1975 although the effect was less prominent, possibly due to H1975’s lower endogenous KIFC1 expression and levels of CA (**Supp. Fig. 4**). We also found that KIFC1 LOF reduced the fitness of H1299 cells treated with DMSO (**Fig.4D**), indicating that KIFC1 is required to compensate for the relatively high basal CA in the H1299 model. KIFC1-perturbation did not reduce the fitness of PC9 or A549 cells with relatively low CA and KIFC1 expression (**Fig.4D**). These results implicate KIFC1 as a specific dependency in LUAD cell lines with high basal CA (H1299, H1975). The lack of sensitization in PC9 suggests this model uses KIFC1-independent mechanisms to cope with CA induction, while the lack of effect in A549 likely reflects their resistance to CA induction.

**Figure 4.**
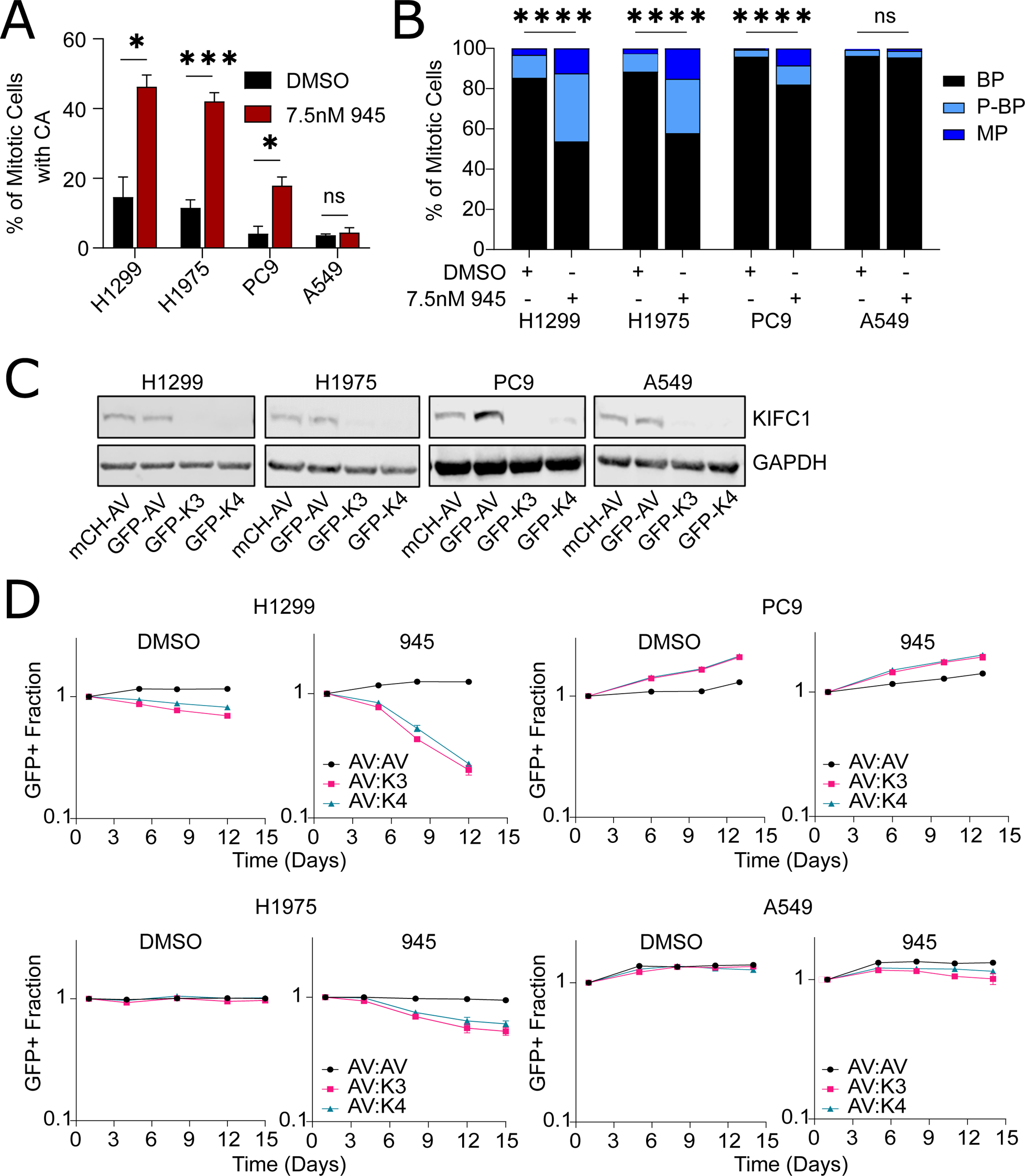
Validation of KIFC1 as a genetic dependency in LUAD models with CA. (A) Potentiation of CA by low-dose 945 treatment (7.5nM) in Cas9+ cell models. Centrosomes were detected using IF for CEP192 and CA was scored as the proportion of cells with greater than two centrosomes. Error bars indicate the mean and SD (N=3), and asterisks indicate p-value significance for t-tests between treatments. (B) Classification of mitotic spindles as bipolar (BP), multipolar (MP), or pseudo-bipolar (P-BP) for each treatment condition. Asterisks indicate p-value significance for a chi-square test. For (A) and (B), data for 3 replicates are pooled and at least 150 total mitotic cells were scored for each treatment condition. (C) Confirmation of CRISPR-mediated loss of KIFC1 expression in LUAD cell lines. Cas9+ cells were transduced with mCherry (mCH) or GFP lentivectors to express sgRNA targeting AAVS1 (AV) or KIFC1 (K3 targeting exon 3 and K4 targeting exon 4). (D) Multicolour competition assays comparing the relative fitness of cells with wildtype (AV) and genetically perturbed *KIFC1* (K3, K4) in DMSO- and 945-treated conditions. mCH-AV cells were mixed 1:1 with GFP-AV, GFP-K3, or GFP-K4 cells and grown over multiple passages. The GFP+ fraction was monitored over time by flow cytometry. Data are representative of two independent experiments (*p<0.05, **p<0.01, ***p<0.001, ****p<0.0001).

### Depletion of KIFC1 impairs cell viability and suppresses centrosome clustering in LUAD cell lines with CA

Although multiple compounds have been reported to inhibit KIFC1[44–46], their weak potencies require treating cells at high doses that preclude attributing treatment-induced phenotypes specifically to KIFC1 inhibition[47]. We investigated the effects of AZ82, the KIFC1 inhibitor reported to have the best specificity[47], on LUAD viability. We observed cytotoxic effects of AZ82 at 7.5uM but they did not correlate with KIFC1 expression (**Supp. Fig. 5**). For example, PC9 exhibited the greatest sensitivity to AZ82 in contrast to the results of our CRISPR screen validation experiments (**Fig.4D; Supp. Fig. 5**). As such, we used two independent siRNAs to knockdown (KD) KIFC1 to investigate its therapeutic potential. Both siRNA yielded effective KD in all 4 models (**Fig.5A-B**). Dose response curves indicated that KIFC1 depletion enhanced sensitivity to 945 in H1299 (**Fig.5C**). A similar albeit less pronounced effect was observed in H1975 and PC9, and KIFC1 KD had no effect on the sensitivity of A549 to 945. We also conducted clonogenic survival assays to measure the effects of KIFC1 KD on cell viability. KIFC1 KD significantly reduced colony survival in H1299 and H1975 when CA was potentiated by 945 treatment, but had no effect in 945-treated PC9 and A549 cells (**Fig.5D**). Depletion of KIFC1 alone had no effect on cell proliferation or clonogenic survival **(Supp. Fig. 6A-B)**.

**Figure 5.**
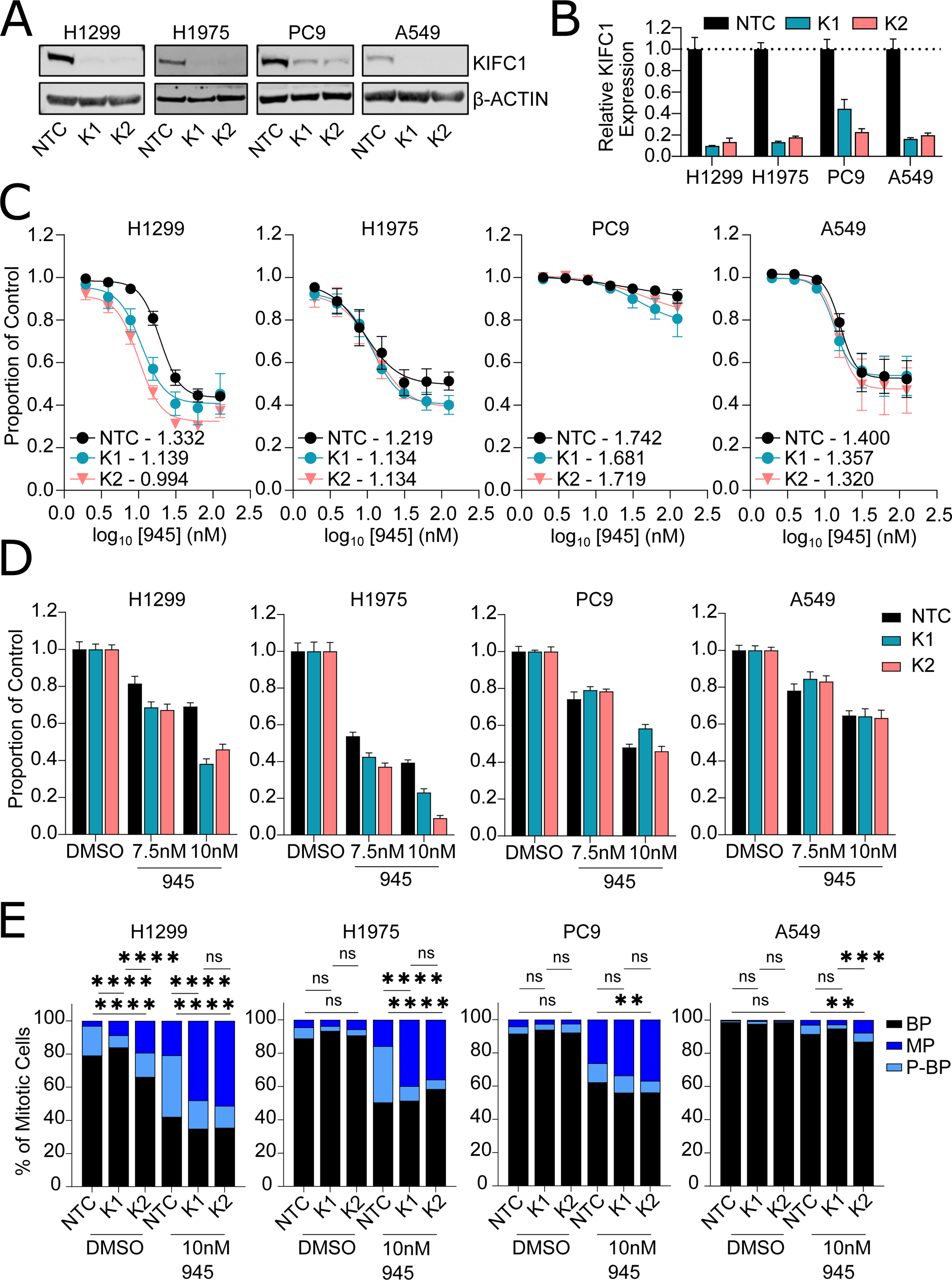
Depletion of KIFC1 sensitizes LUAD models with high basal CA to potentiation of CA. (A,B) Confirmation of siRNA-mediated KIFC1 knockdown by western blotting (A) and RT-qPCR (B). Relative quantification (RQ; 2^-ΔΔCt^) values were calculated using the ΔΔCt method and error bars indicate the minimum and maximum RQ for 3 technical replicates. RNA and protein were collected 24hr post transfection at the time that cells were seeded for functional assays. Representative knockdowns are shown. NTC = non-targeting control. K1, K2 indicate independent siRNA targeting KIFC1. (C) Dose response assays for 945 treatment in cells with and without KIFC1 knockdown. Cell viability at each dose was normalized to that for DMSO-treated controls. Each curve represents mean viability data for 3-5 biological replicates. Error bars indicate SEM. Numbers indicate area under the curve. (D) Clonogenic survival assays. Data are representative of 2-3 biological replicates per model. Colony survival was normalized to that observed in DMSO-treated conditions for each siRNA. Error bars indicate SEM of 6 technical replicates. (E) Classification of mitotic spindles as bipolar (BP), multipolar (MP), or pseudo-bipolar (P-BP). Data are pooled for 3-4 replicate experiments and at least 150 total mitotic cells were scored for each treatment condition. Asterisks indicate p-value significance for chi-square tests between NTC and K1 or K2 siRNA for each treatment condition (*p<0.05, **p<0.01, ***p<0.001, ****p<0.0001).

CA can lead to the formation of multipolar spindles during mitosis (**Fig.1D-E**) and KIFC1 counters their deleterious effects by clustering extra centrosomes to promote viable chromosome segregation and cell division. We suspected that the dependence of LUAD cells with CA on KIFC1 was due to its role in centrosome clustering, so we evaluated the effects of KIFC1 KD on spindle polarity (**Fig.5E**). KIFC1 depletion in H1299 led to a decrease in centrosome clustering evident by a reduced proportion of mitotic cells with pseudo-bipolar spindles and an increased proportion of cells with multipolar spindles. This effect was more pronounced when CA was potentiated with 945 treatment. KIFC1 KD did not affect centrosome clustering in H1975, PC9, or A549 cells treated with DMSO. Like H1299, 945 treatment reduced pseudo-bipolar and increased multipolar spindles in H1975 (**Fig.5E**). KD of KIFC1 in PC9 and A549 increased the proportion of multipolar spindles in 945-treated cells but this phenotype was not reproduced with both siRNAs and the magnitude of change was less than that observed for H1299 and H1975 (**Fig.5E**). These data suggest that KIFC1 promotes survival of LUAD cells with CA by clustering extra centrosomes into pseudo-bipolar spindles to avoid the lethal consequences of cell divisions with multipolar spindles.

## DISCUSSION

Centrosome amplification (CA) is a recurrent feature of many malignancies that promotes invasiveness and genomic instability (GIN), and consequently, cancer development and progression[12, 13, 42, 43, 48]. Not surprisingly, CA is associated with poor prognosis in many cancer types[17, 21, 29, 49]. Despite the prevalence of CA in lung tumors[6, 7, 17], the vulnerabilities imposed by CA in LC and its therapeutic potential have not been defined. By integrating a CRISPR/Cas9 screen, genomic analyses of LUAD tumors, and functional studies in multiple cell lines, we identified and validated *KIFC1* as a dependency of LUAD with supernumerary centrosomes. Kinesin family member C1 (KIFC1 or HSET) belongs to the kinesin-14 family of minus-end directed motor proteins that facilitate organization of the mitotic spindle[33]. Both the tail and motor domains of KIFC1 bind to microtubules, allowing it to crosslink microtubules and slide them apart or together to organize spindle poles[50, 51]. Although KIFC1 is not essential in non-malignant cells, it is indispensable in some cancer cell lines[52, 53]. Studies of multipolar divisions in cells with CA have demonstrated that KIFC1 clusters extra centrosomes into spindles with pseudo-bipolar arrangements[42, 52, 54, 55]. This action is thought to occur by KIFC1 ratcheting microtubules emanating from different centrosomes to group them together[54].

Our discovery of KIFC1 as a putative vulnerability in LC is consistent with reports in breast, prostate, ovarian, liver, and renal cancers[45, 52, 56–58]. Liu and colleagues investigated KIFC1 using functional studies in 3 NSCLC lines and reported that KIFC1 depletion reduced viability and induced G2/M arrest and p21 expression but they did not evaluate centrosomes[35]. However, they reported that KIFC1 KD reduced viability to the greatest extent in H1299 cells, which we found to have relatively high basal CA among the LUAD models we assessed. Advancing on correlative studies that did not consider centrosome number, our unique CA-focused approach nominated KIFC1 as a dependency specifically in LUAD models with high basal levels of CA, likely due to its established role in mitigating CA-associated mitotic stress via centrosome clustering[45, 52, 56–58]. Patel *et al*. identified KIFC1 as a putative vulnerability in breast cancers based on integrative genomic analyses of tissues and cell lines. Similar to our finding, this group showed that KIFC1 was preferentially essential in triple-negative breast cancer models with CA[57]. Although the CA20 gene expression signature is only a surrogate for CA, our assessment of KIFC1 dependency and CA20 in the DepMap database[20] further suggested that KIFC1 is a CA-specific dependency in LUAD and possibly SCLC. KIFC1’s classification as a strongly selective dependency, but not a common essential gene, in DepMap likely indicates that a small fraction of cancer models with high basal CA are dependent on KIFC1.

Liu *et al*. also reported that KIFC1 was overexpressed in NSCLC compared to non-malignant lung tissues and that high expression was associated with worse prognosis, which is consistent with findings from other LC studies[35–37, 59, 60] and reports in other malignancies[34, 36, 61–65]. Our analyses of multiple LC cohorts confirmed associations of KIFC1 with clinical factors including survival and stage as well as CA20. For the first time, we found that KIFC1 expression was elevated in LUAD from patients with a smoking history. This observation was reproducible across multiple cohorts, raising the possibility that tobacco smoke leads to increased KIFC1 expression as a consequence of inducing CA[66]. Celik *et al*. recently reported that KIFC1 was overexpressed in 59% of lung tumors and that 96% of these exhibited hypomethylation of KIFC1[59]. This suggests that recurrent overexpression of KIFC1 in LC is driven by epigenetic mechanisms, but this remains to be validated in additional cohorts. An integrative pan-cancer study by Wu *et al*. found that KIFC1 was rarely perturbed at the genetic level in LUAD and other tumor types, suggesting genetic mechanisms are unlikely to account for KIFC1 overexpression[34]. Further studies are needed to decipher the mechanisms driving KIFC1 overexpression but ample evidence indicates that KIFC1 expression is a robust prognostic biomarker in LC.

Targeting kinesins that regulate spindle assembly in cancer cells is a promising therapeutic approach particularly in genomically unstable tumors[67–69]. Inhibitors of KIF18A and Eg5 (encoded by *KIF11*) are being investigated in clinical trials[68, 70]. Our work nominates KIFC1 as a therapeutic target in LC, with CA as a putative predictive biomarker. Our finding that KIFC1 expression was significantly higher in LUAD lacking common driver mutations and in smokers compared to former and never smokers suggests that KIFC1-targeted therapy could be useful in these currently underserved patient populations. Encouragingly, KIFC1 KD does not affect proliferation or survival in non-malignant cells *in vitro*[52, 57, 71], but studies of KIFC1-targeted therapy in animal models are required to confirm its therapeutic index. None of the KIFC1 inhibitors described to date (e.g. AZ82, CW069, SR17852, and KAA[44–46, 72]) have progressed to clinical studies, which may be attributable to poor potencies, specificities, and/or bioavailability[47, 73]. Thus, development of clinical-grade KIFC1 inhibitors is warranted, especially given their potential utility in diverse cancers. Our analyses of CA and the CA20 gene signatures suggest that various opportunities exist to measure CA as a potential biomarker. As such, validation of these candidate response-predictive features could facilitate the translation of KIFC1-targeted therapy by informing rational patient selection.

Our work and that of others indicates that cultured cancer cells exhibit less CA than malignant cells in tumor tissues[74]. Accordingly, we treated LC cells with low dose 945[30] to potentiate CA and better mimic clinical tumors and observed that KIFC1 LOF sensitized cells to CA. This observation was consistent with a study that reported that combining KIFC1 KD with potentiation of CA by cisplatin enhanced cytotoxicity in TNBC cells[57]. Together, these results suggest that KIFC1 inhibition could synergize with chemotherapy and/or irradiation which induce CA and/or multipolar spindles[11, 75, 76]. Supporting this concept, a recent study showed that KIFC1 KD sensitized LC to radiotherapy, though CA was not measured[71]. Further studies to define rational and effective KIFC1 inhibitor combinations are warranted, but our work nominates 945 as a promising partner for KIFC1-targeted therapy. Confirming whether 945 treatment enhances CA in tumor tissues from treated patients would provide additional evidence to support this strategy. Since 945 does not induce CA in non-malignant cells, this combination could be clinically tractable[30, 77, 78].

The recurrence of CA and KIFC1 overexpression in lung tumors, and the dependence of LC with CA on KIFC1 emphasize its therapeutic potential[33, 34]. We propose that KIFC1-targeted therapy could provide a novel strategy for potentiating GIN to lethal levels by forcing LC to divide with multipolar spindles, and that CA and KIFC1 expression could serve as biomarkers to guide effective use. We believe the efficacy of KIFC1-targeted therapy will be greatest when combined with treatments that exacerbate CA in lung tumors, and suggest that developing clinical-grade KIFC1 inhibitors and identifying and evaluating rational therapeutic combinations is warranted.

## Acknowledgements

We thank the UHN Therapeutics group for providing CFI-400945 for our studies, TCGA for tumor genomics data, and the late Dr. Adi Gazdar for sharing LUAD cell lines. This work was supported by St. Michael’s Hospital, the Canada Research Chairs program, the Canadian Foundation for Innovation, and the Canadian Institutes of Health Research (CIHR). C.Z. was supported by a CIHR scholarship.

## Author Contributions

CZ, BW, and KLT were responsible for the study conception and design, and manuscript preparation. CZ and BW conducted all functional experiments with assistance from YFW and KLT. CZ, BW, and KLT performed data analyses. NAP and MST provided PDX tissue samples. AE performed IHC experiments. RC and ISB provided reagents and assisted with custom library design and synthesis. CDCO contributed to microscopy data acquisition and processing. WLL provided clinical data for the BCCA dataset (GSE75037). MRB, TWM and DWC provided CFI-400945 and other essential reagents. All authors made significant contributions and critically reviewed the manuscript.

## Data Availability

Datasets analyzed are available from public sources as described, and CRISPR screen data are available from the corresponding author upon reasonable request.

---

**Supplementary Figure 1. Technical validation of H1299 CRISPR/Cas9 screen.** (A) Confirmation of Cas9 editing efficiency in H1299-Cas9 cells. Cas9+ cells were transduced with either a BFP-GFP-empty vector or a BFP-GFP-sgGFP vector, and GFP expression was quantified by flow cytometry 48hr later. The percentage of BFP+GFP+ cells is indicated in Q2. (B) Potentiation of centrosome amplification in H1299-Cas9 cells by treatment with low doses of 945. Centrosomes were scored based on immunofluorescence for CEP192. Error bars indicate mean and SD for two replicates. (C) Growth curves of DMSO and 945-treated cell populations during the CRISPR screen. Error bars indicate the mean and SD for the 3 technical replicates. (D) Identification of essential genes in H1299 cells from the CRISPR screen. Normalized Z-scores for target genes in DMSO-treated cells at screen end point relative to screen onset (T0) are shown. Z-scores represent the relative abundance of sgRNAs targeting each gene in the cell populations. Negative Z-scores indicate genes targeted by sgRNAs that were significantly depleted in DMSO-treated cells at the end of the screen. (E) Abundance of sgRNA targeting *MYC* and *NRAS* at screen onset (T0) compared to endpoint (after 17 doublings).

**Supplementary Figure 2. Association of KIFC1 expression with clinical features in additional LUAD cohorts.** A) KIFC1 expression in LUAD from males and females in the TCGA dataset. B) KIFC1 expression in the TCGA cohort segregated by mutation status for common oncogenic drivers in LUAD: *EGFR, KRAS, MET, ROS1, RET, ALK, NRAS* and *BRAF*. In 4 additional LUAD datasets from the gene expression omnibus, KIFC1 expression was investigated in relation to smoking history (B), sex (C), stage (D), CA20 score (E), and 5-year survival (F). Asterisks indicate significance for Mann Whitney or Kruskal-Wallis tests. Error bars indicate mean and SD. NS = never smokers, CS = current smokers, ES = ever smokers, FS = former smoker. Pearson’s correlation coefficients are indicated for associations between KIFC1 expression and CA20 scores. *Note that only 19/20 genes in the CA20 signature were evaluable in the GSE68465. Association between KIFC1 expression and 5-year survival was assessed in patients whose tumors exhibited KIFC1 expression ranking in the top and bottom tertiles of expression. Asterisks indicate significance for log-rank tests on the Kaplan-Meier survival curves (*p<0.05, **p<0.01, ***p<0.001, ****p<0.0001).

**Supplementary Figure 3. Association of KIFC1 expression with clinical features in lung squamous cell carcinoma (LUSC) patients.** KIFC1 expression was analyzed in the TCGA LUSC cohort to investigate associations with malignancy (A), smoking history (B), sex (C), and stage (D). Asterisks indicate significance for Mann Whitney or Kruskal-Wallis tests (****p<0.0001). Error bars indicate mean and SD. NS = never smokers, CS = current smokers, FS < 15 = former smoker for fewer than 15 years, FS > 15 = former smoker for over 15 years. (E) Association between KIFC1 expression and 5-year survival was assessed in patients whose tumors exhibited KIFC1 expression ranking in the top and bottom tertiles of expression. The survival association was investigated using a log-rank test on the Kaplan-Meier survival curves. (F) Correlation between KIFC1 mRNA expression and CA20 scores. Correlation coefficient is that for Pearson’s correlation.

**Supplementary Figure 4. Centrosome amplification and KIFC1 expression in LUAD cell lines.** (A) Western blot for KIFC1 expression in 14 LUAD lines. (B) Quantification of protein expression in (A). KIFC1 levels were normalized to GAPDH. (C) Proportion of cells with centrosome amplification in 11 LUAD lines. Centrosomes were identified by immunofluorescence staining for CEP192. (D) Correlation between KIFC1 dependency and CA20 in LUSC (D) and SCLC (E) cell lines from the CCLE. KIFC1 gene effect scores were accessed from the DepMap database and negative scores indicate genetic dependency. CA20 scores were obtained from the de Almeida 2019 study. The correlation coefficients indicated were calculated using Pearson’s correlation analyses.

**Supplementary Figure 5. Cytotoxicity of AZ82 in LUAD cell lines.** LUAD cells were treated for 5 days with DMSO or AZ82 at two doses, 3.5uM (A) and 7.5uM (B). Surviving cells were fixed, stained with SRB, and solubilized for colorimetric readouts on a spectrophotometer. Values indicate viability in AZ82-treated cells relative to DMSO controls. Error bars indicate SD for replicate experiments.

**Supplementary Figure 6.** Effect of KIFC1 knockdown on proliferation and clonogenic survival. A) Cell proliferation was measured using the ATP-Lite assay over 5 days. Values indicate luminescence signals normalized to Day 0. Growth curves represent data pooled from 4-6 replicate experiments and error bars indicate SEM. B) Raw colony numbers for each siRNA condition are plotted for the DMSO-treated cells from Fig.5. Colonies were grown for 11-14 days. Colony numbers in cells transfected with NTC and K1 or K2 siRNA were compared using t-tests. Data represent 3-5 experimental replicates. Error bars indicate SEM.

