## Supplementary Tables and Figures for "Identification of KIFC1 as a putative vulnerability in lung cancers with centrosome amplification"

#### SUPPLEMENTARY MATERIAL

**Supplementary Table 1. Sequences for the custom sgRNA library targeting druggable genes.**
<Excel file attached>

**Supplementary Table 2. sgRNA and qPCR primer sequences.**

| **sgRNA** | **Sequence** |
|---|---|
| **sgAAVS1 (AV)** | GGGGGCCACTAGGGACAGGAT |
| **sgKIFC1 (K3)** | CAGACACAAGGCCAGACCAC |
| **sgKIFC1 (K4)** | GCAATAGCTGTGGAACACCG |
| **qPCR primer** | **Sequence** |
| **hs.HPRT1.F** | CCTGGCGTCGTGATTAGTGAT |
| **hs.HPRT1.R** | AGACGTTCAGTCCTGTCCATAA |
| **hs.KIFC1.F** | CCTGGAGCCTGAGAAGAAACG |
| **hs.KIFC1.R** | GGTAATTTTGGTCGTTGCACCC |

**Supplementary Table 3. Top hits identified by the CA-sensitization screen in H1299 cells.**

| Gene | Norm Z score | pval_ synth | Rank_ synth | FDR_ synth | Correlation with CA20 (TCGA LUAD) |
|---|---|---|---|---|---|
| **KIFC1** | -3.07 | 0.00107 | 15 | 0.236 | 0.8990 |
| **ELAVL1** | -3.33 | 0.000435 | 10 | 0.144 | 0.4533 |
| **PTGES3** | -3.73 | 9.58E-05 | 3 | 0.106 | 0.4063 |
| **CFL1** | -3.49 | 0.000245 | 6 | 0.136 | 0.3728 |
| **POLE4** | -3.18 | 0.000737 | 11 | 0.223 | 0.2034 |
| **NDUFC2** | -4.49 | 3.65E-06 | 1 | 0.0121 | 0.1779 |
| **WEE1** | -4.38 | 5.93E-06 | 2 | 0.00985 | 0.0860 |
| **PHYH** | -3.06 | 0.00111 | 16 | 0.23 | 0.0850 |
| **BRPF3** | -3.41 | 0.000326 | 8 | 0.135 | 0.0435 |
| **APRT** | -3.5 | 0.000234 | 4 | 0.194 | -0.0283 |
| **RBBP9** | -3.42 | 0.000318 | 7 | 0.151 | -0.1272 |
| **NOX1** | -3.11 | 0.000937 | 13 | 0.239 | -0.1704 |
| **RXRB** | -3.08 | 0.00103 | 14 | 0.244 | -0.1737 |
| **SMG6** | -3.49 | 0.000244 | 5 | 0.162 | -0.3455 |
| **AXIN2** | -3.18 | 0.000743 | 12 | 0.206 | -0.3931 |
| **CYB5R3** | -3.33 | 0.000427 | 9 | 0.158 | -0.4414 |

Supp.Fig.1

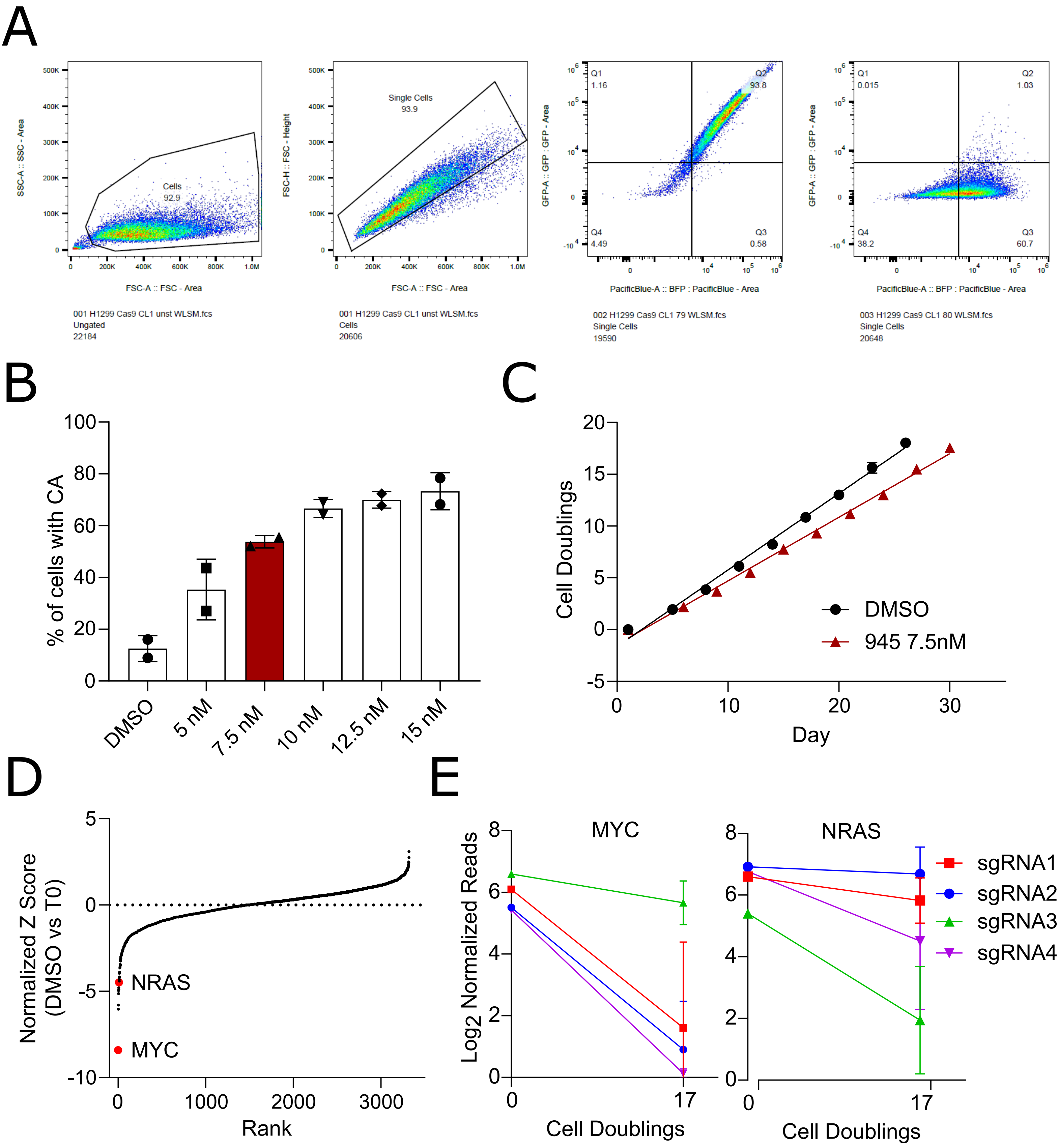
A
Cells
92.9
SSC-A :: SSC - Area
FSC-A :: FSC - Area
001 H1299 Cas9 CL1 unst WLSM.fcs
Ungated
22184
Single Cells
93.9
FSC-H :: FSC - Height
FSC-A :: FSC - Area
001 H1299 Cas9 CL1 unst WLSM.fcs
Cells
20606
Q1
1.16
Q2
93.8
Q4
4.49
Q3
0.58
GFP-A :: GFP : GFP - Area
PacificBlue-A :: BFP : PacificBlue - Area
002 H1299 Cas9 CL1 79 WLSM.fcs
Single Cells
19590
Q1
0.015
Q2
1.03
Q4
38.2
Q3
60.7
GFP-A :: GFP : GFP - Area
PacificBlue-A :: BFP : PacificBlue - Area
003 H1299 Cas9 CL1 80 WLSM.fcs
Single Cells
20648
B
% of cells with CA
DMSO
5 nM
7.5 nM
10 nM
12.5 nM
15 nM
C
Cell Doublings
Day
DMSO
945 7.5nM
D
Normalized Z Score
(DMSO vs T0)
NRAS
MYC
Rank
E
Log2 Normalized Reads
MYC
NRAS
Cell Doublings
Cell Doublings
sgRNA1
sgRNA2
sgRNA3
sgRNA4

### Supp.Fig.2

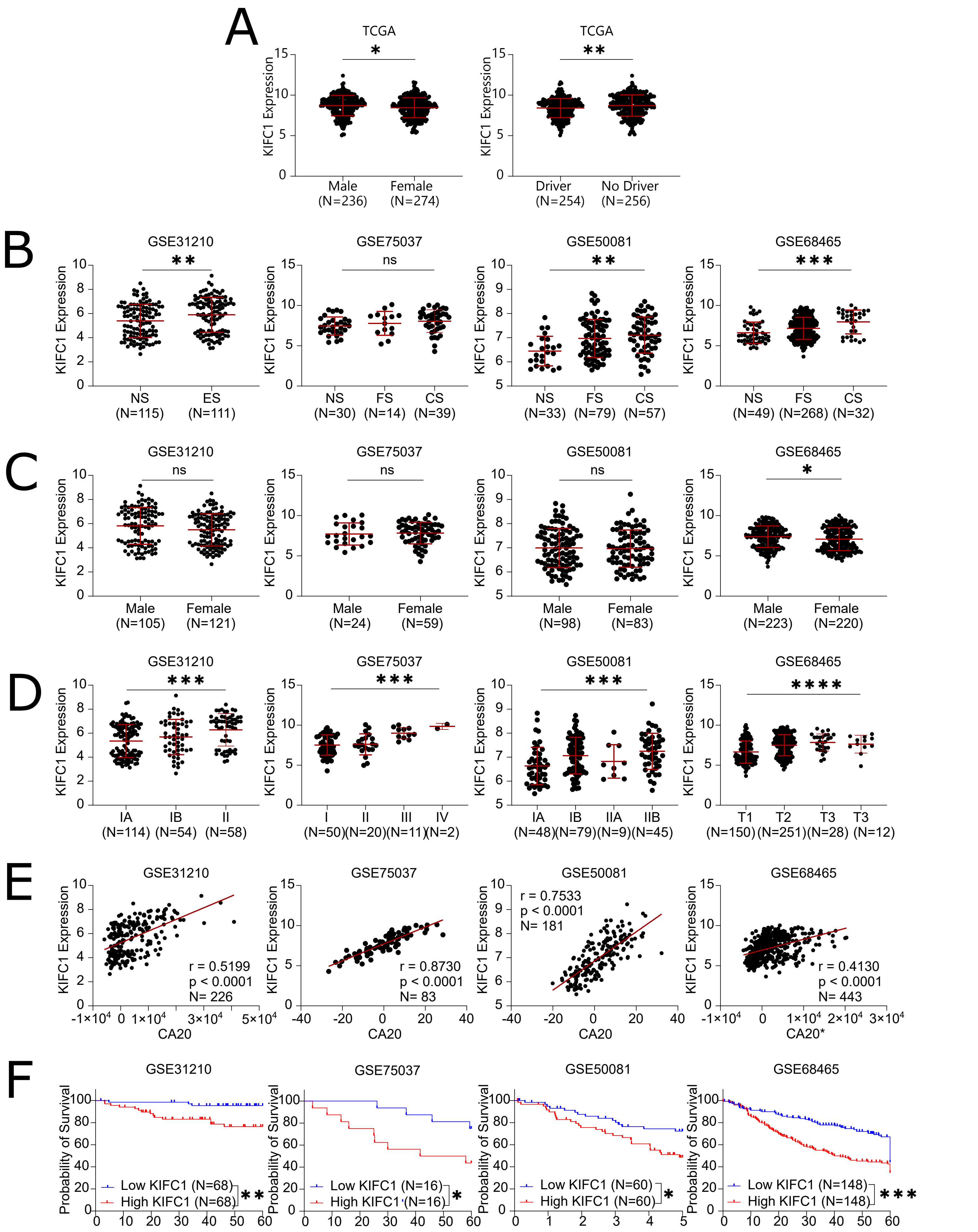
A
TCGA
KIFC1 Expression
*
Male (N=236)
Female (N=274)
TCGA
**
Driver (N=254)
No Driver (N=256)
B
GSE31210
**
NS (N=115)
ES (N=111)
GSE75037
ns
NS (N=30)
FS (N=14)
CS (N=39)
GSE50081
**
NS (N=33)
FS (N=79)
CS (N=57)
GSE68465
***
NS (N=49)
FS (N=268)
CS (N=32)
C
GSE31210
ns
Male (N=105)
Female (N=121)
GSE75037
ns
Male (N=24)
Female (N=59)
GSE50081
ns
Male (N=98)
Female (N=83)
GSE68465
*
Male (N=223)
Female (N=220)
D
GSE31210
***
IA (N=114)
IB (N=54)
II (N=58)
GSE75037
***
I (N=50)
II (N=20)
III (N=11)
IV (N=2)
GSE50081
***
IA (N=48)
IB (N=79)
IIA (N=9)
IIB (N=45)
GSE68465
****
T1 (N=150)
T2 (N=251)
T3 (N=28)
T3 (N=12)
E
GSE31210
r = 0.5199
p < 0.0001
N= 226
CA20
GSE75037
r = 0.8730
p < 0.0001
N= 83
CA20
GSE50081
r = 0.7533
p < 0.0001
N= 181
CA20
GSE68465
r = 0.4130
p < 0.0001
N= 443
CA20*
F
Probability of Survival
GSE31210
Low KIFC1 (N=68)
High KIFC1 (N=68)
**
GSE75037
Low KIFC1 (N=16)
High KIFC1 (N=16)
*
GSE50081
Low KIFC1 (N=60)
High KIFC1 (N=60)
*
GSE68465
Low KIFC1 (N=148)
High KIFC1 (N=148)
***

Supp.Fig.3

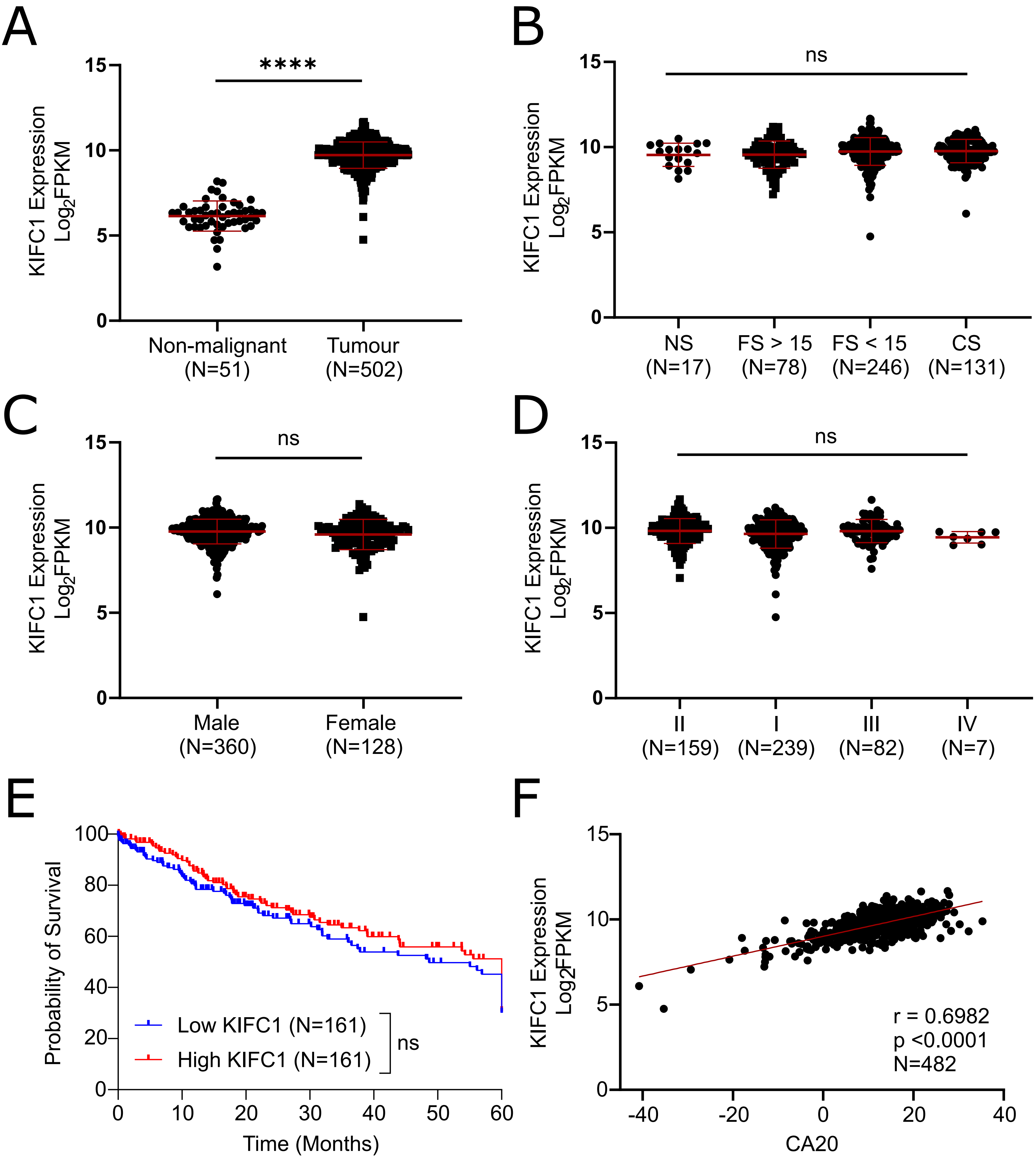
A
****
KIFC1 Expression Log2FPKM
15
10
5
0
Non-malignant (N=51)
Tumour (N=502)
B
ns
KIFC1 Expression Log2FPKM
15
10
5
0
NS (N=17)
FS > 15 (N=78)
FS < 15 (N=246)
CS (N=131)
C
ns
KIFC1 Expression Log2FPKM
15
10
5
0
Male (N=360)
Female (N=128)
D
ns
KIFC1 Expression Log2FPKM
15
10
5
0
II (N=159)
I (N=239)
III (N=82)
IV (N=7)
E
Probability of Survival
100
80
60
40
20
0
Low KIFC1 (N=161)
High KIFC1 (N=161)
ns
0
10
20
30
40
50
60
Time (Months)
F
KIFC1 Expression Log2FPKM
15
10
5
0
r = 0.6982
p <0.0001
N=482
-40
-20
0
20
40
CA20

### Supp.Fig.4

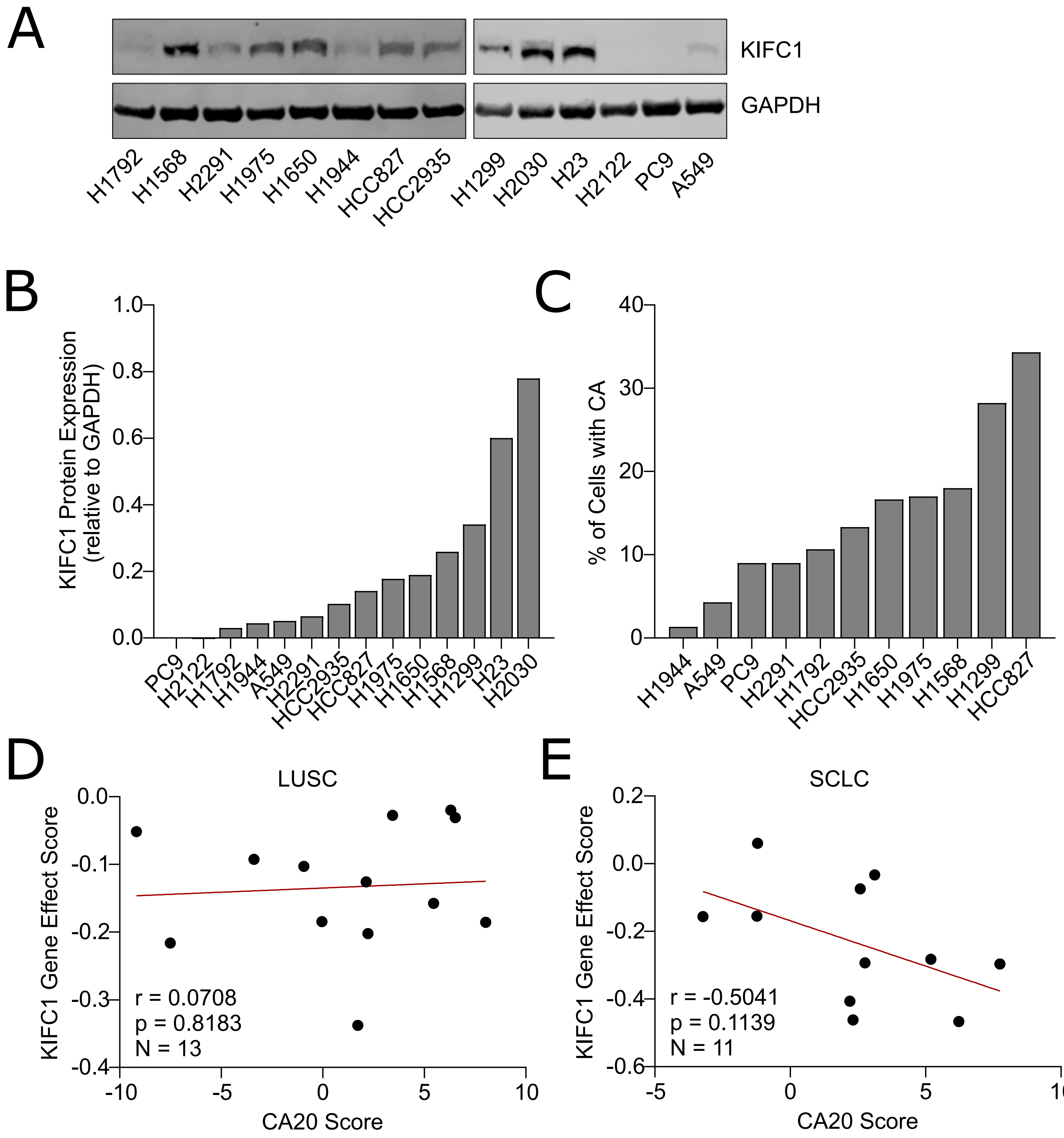
A
KIFC1
GAPDH
H1792
H1568
H2291
H1975
H1650
H1944
HCC827
HCC2935
H1299
H2030
H23
H2122
PC9
A549
B
KIFC1 Protein Expression (relative to GAPDH)
1.0
0.8
0.6
0.4
0.2
0.0
PC9
H2122
H1792
H1944
A549
H2291
HCC2935
HCC827
H1975
H1650
H1568
H1299
H23
H2030
C
% of Cells with CA
40
30
20
10
0
H1944
A549
PC9
H2291
H1792
HCC2935
H1650
H1975
H1568
H1299
HCC827
D
LUSC
KIFC1 Gene Effect Score
0.0
-0.1
-0.2
-0.3
-0.4
r = 0.0708
p = 0.8183
N = 13
-10
-5
0
5
10
CA20 Score
E
SCLC
KIFC1 Gene Effect Score
0.2
0.0
-0.2
-0.4
-0.6
r = -0.5041
p = 0.1139
N = 11
-5
0
5
10
CA20 Score

### Supp.Fig.5

A

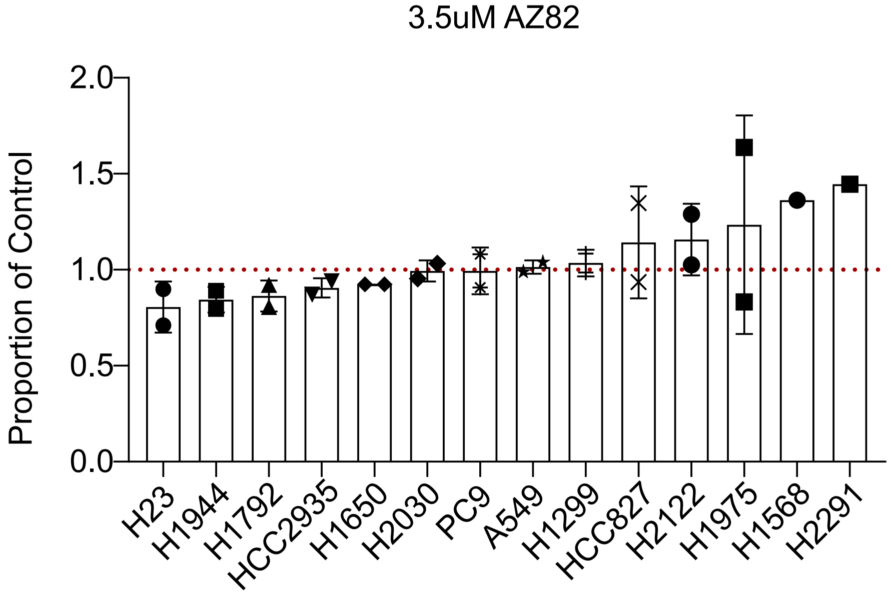
3.5uM AZ82
Proportion of Control
2.0
1.5
1.0
0.5
0.0
H23
H1944
H1792
HCC2935
H1650
H2030
PC9
A549
H1299
HCC827
H2122
H1975
H1568
H2291

B

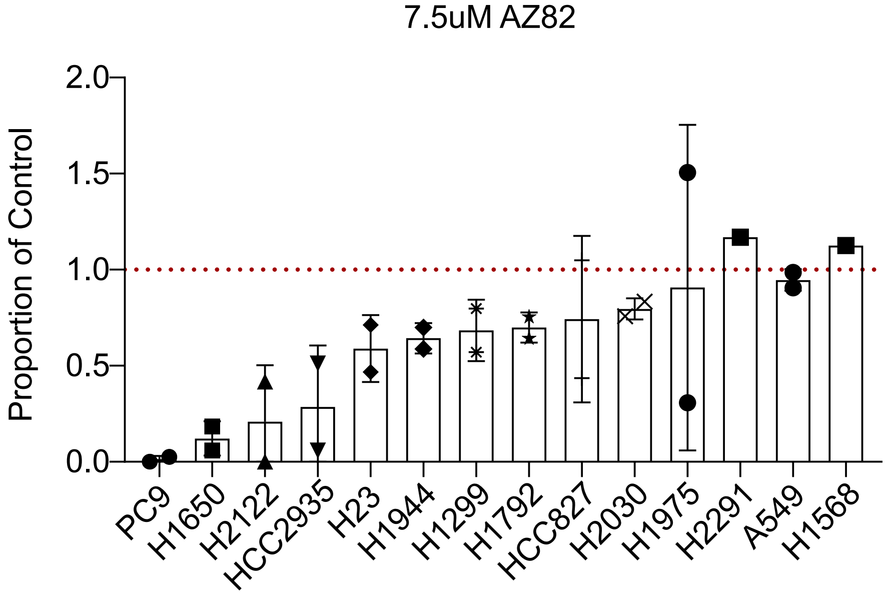
7.5uM AZ82
Proportion of Control
2.0
1.5
1.0
0.5
0.0
PC9
H1650
H2122
HCC2935
H23
H1944
H1299
H1792
HCC827
H2030
H1975
H2291
A549
H1568

### Supp.Fig.6

A

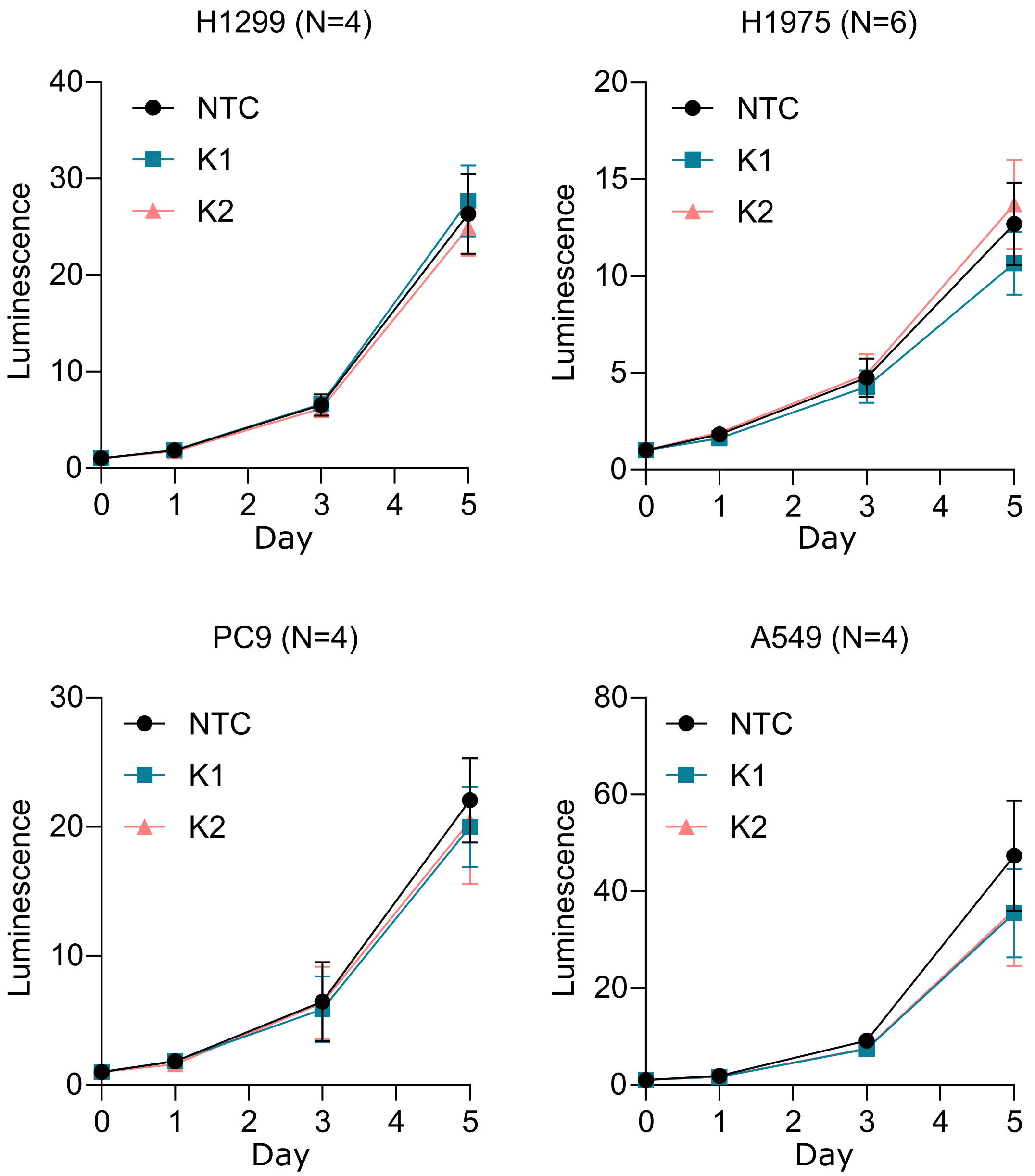
H1299 (N=4)
H1975 (N=6)
PC9 (N=4)
A549 (N=4)
NTC
K1
K2
Luminescence
Day
0
1
2
3
4
5
10
15
20
30
40
60
80

B

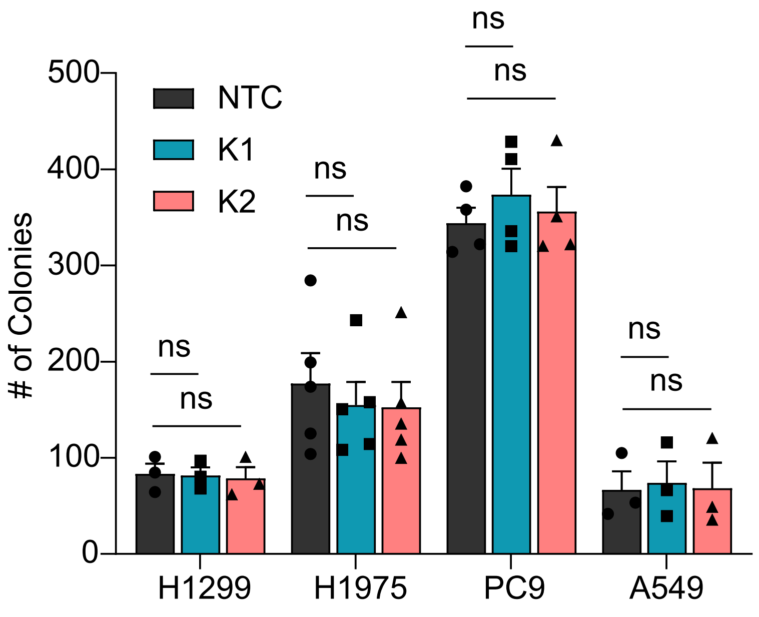
NTC
K1
K2
### of Colonies
ns
0
100
200
300
400
500
H1299
H1975
PC9
A549
